# Bumblebees learn to use peripheral taste to predict the presence of nectar in flowers

**DOI:** 10.1101/2025.03.11.642579

**Authors:** Michael J.M. Harrap, Rachel H. Parkinson, Hannah Jones, Geraldine A. Wright

## Abstract

Learning cues such as tastes associated with palatable food is an important mechanism animals have for foraging optimally. Insects can use gustatory receptor neurons (GRNs) in their mouthparts to detect nutrients and toxins, but they also taste compounds using sensilla on peripheral organs such as their antennae. Bees are adept at learning to associate floral traits with the presence of nectar rewards, but few studies have examined how they incorporate gustatory information from their antennae with rewards. Here, we characterize the ability of adult worker bumblebees (Bombus terrestris) to taste sugar, salt, and bitter compounds using their antennae and then tested whether they could use this sensory information to associate it with food. We show that bumblebees have antennal GRNs sensitive to sugars, salts, and bitter compounds and that they can use surface chemistry differences detected by their periphery to learn about the presence or absence of flower rewards in a free-flight assay. Naïve bumblebees showed no instinctual preferences toward or against any surface chemistry tested. Bumblebees performed best when sucrose surface cues were associated with sucrose reward, but they could learn to associate any cue with the presence or absence of sucrose solution. Interestingly, the bees found it more difficult to associate quinine surface chemistry with the presence of reward than its absence. These results indicate that bees have the potential to learn to associate another floral trait – chemicals on the surfaces of petals - with the quality of floral rewards.

**Summary statement:** Behavioural experiments and electrophysiological recordings show bumblebees can detect peripheral taste cues on surfaces of artificial flowers, including bitter toxins, and learn to use these to predict rewards.

## Introduction

The sense of taste guides the intake of food by sensing nutrients and makes it possible for an animal avoid potential toxins (Beauchamp and Jiang, 2015; Beauchamp, 2016). Gustatory information directly actives the neural circuits that control instinctual food consumption and rejection, but it also informs the brain of the value of food and can therefore influence learning and memory (Thoma et al., 2016). Animals have taste structures (e.g. buds in vertebrates or sensilla in insects) located on their mouths at the point of food consumption. In addition to the sensilla that house gustatory neurons (GRNs) on the mouthparts, insects and arthropods can also detect tastants peripherally on their antennae, tarsi, and even ovipositors (Bestea et al., 2021; Meunier et al., 2003; Mitchell et al., 1999; Shrestha and Lee, 2023; Whitehead and Larsen, 1976). Insect GRNs spike in response tastants such as sugars and other nutrients, salt, water, and potential ‘bitter’ compounds (Dethier, 1976). The relative levels of activity of the GRNs inform the brain of food value and guide behaviour (Wright, 2016). For example, stimulation of gustatory sensilla with sucrose in all locations evokes instinctual feeding reflexes (Dethier, 1976; Guerrieri and d’Ettorre, 2010; Kuwabara, 1957; Mommaerts et al., 2013; Villagómez et al., 2024; Zhang et al., 2010), whereas stimulation with bitter compounds provokes aversion or food rejection (Bernays and Chapman, 2000; Chapman and Bernays, 1989; de Brito Sanchez et al., 2005; Hostachy et al., 2019; Ma et al., 2016; Matsumoto et al., 2015; Nicholls et al., 2019). Several studies indicate that stimulation of insect mouthparts sensilla is important for learning and memory (Chabaud et al., 2006; Guerrieri and d’Ettorre, 2010; Hartlieb et al., 1999; Laloi et al., 1999; Matsumoto et al., 2015; Nicholls et al., 2019; Riveros and Gronenberg, 2009; Simcock et al., 2018), but the extent to which peripheral gustatory sensilla contribute to sensory representations that inform learning and memory has been rarely studied.

Several advantages are conveyed to insects by having gustatory sensilla located peripherally (i.e. antennae). Peripheral gustation increases an animal’s sensitivity to detect small quantities of food and to orient mouthparts towards nutrients (Brockmann et al., 2018; Corfas et al., 2019; Dethier, 1976; Mommaerts et al., 2013; Titova et al., 2023). Insects can also detect potential toxins in food sources using antennae or tarsi, and this allows them to completely avoid intoxication by ingesting toxins (Ayestaran et al., 2010; Barlow et al., 2017; Tiedeken et al., 2014; Villalona et al., 2020; White and Chapman, 1990) or loss of fitness through oviposition mistakes (Newland, 1998; Newland and Yates, 2008; Rogers and Newland, 2000). Being able to associate tastes sensed at the periphery could also be advantageous. For example, bees forage for floral nectar as their main source of carbohydrates (Ansaloni et al., 2025; Heinrich, 2004). Nectar is often concealed in floral corollas: detecting its presence via oral taste is only possible when the bee tries to consume nectar by inserting its mouthparts into the flower. To save time sampling flowers that are unrewarding, bees have evolved instinctual preferences towards various floral traits that are consistently associated with rewards (Dunlap et al., 2019; Dyer et al., 2006; Raine and Chittka, 2007). They are also adept at learning to associate floral traits with the presence, quality and absence of rewards (Dyer et al., 2008; Dyer and Chittka, 2004a; Clarke et al., 2013; Harrap et al., 2021). Learning floral traits allows bees to adjust foraging behaviour dynamically to favour the most rewarding flowers they encounter. This is true even in circumstances where they must learn to go against their innate preferences for floral traits if flower species presenting such traits are more rewarding (e.g. Raine and Chittka, 2008; Whitney et al., 2008; Harrap et al., 2021). If taste cues on floral surfaces were detected by antennae or tarsi, they could improve the efficiency of foraging (Chittka et al., 2009; Raine and Chittka, 2007; Raine and Chittka, 2008), as they would contribute to the multimodal representations of flowers (Kunze and Gumbert, 2001; Leonard et al., 2011a; Leonard et al., 2011b; Leonard et al., 2012).

Many plant species defend flowers from being eaten by herbivores using toxic chemical compounds (Barlow et al., 2017; Manson et al., 2012; Mithöfer and Boland, 2012; Pacini and Hesse, 2005; Palmer-Young et al., 2019; Stevenson, 2020; Wright et al., 2013). It is possible that bees could detect these compounds in floral surfaces and use them as taste cues to learn about the presence, absence or quality of rewards offered by flowers. However, the sensitivity of antennal GRNs in bees to potential toxins is poorly understood. Honeybees have GRNs that respond to sugars such as sucrose applied to the mouthparts sensilla or to their antennae and tarsi (de Brito Sanchez et al., 2005; de Brito Sanchez et al., 2008; de Brito Sanchez et al., 2014; Jung et al., 2015). Honeybees can learn to associate odour stimuli with sucrose solution applied to the antennal sensilla (Bitterman et al., 1983; de Brito Sanchez et al., 2008), but this form of learning does not produce a long-term memory of the odour (Wright et al., 2005b). Like honeybees, bumblebees have sugar sensitive GRNs in their mouthparts (Kessler et al., 2015; Miriyala et al., 2018; Parkinson et al., 2022). GRNs that spike in response to bitter compounds have not been identified at honeybee and bumblebee mouthparts, and honeybee peripheries. These studies identify bitter compounds inhibit the bees sugar sensing GRNs, allowing ‘indirect’ taste of bitter compounds by modulation of sweet taste (de Brito Sanchez et al., 2005; de Brito Sanchez et al., 2008; de Brito Sanchez et al., 2014; Miriyala et al., 2018; Parkinson et al., 2022). However, the responses of GRNs of bumblebee antennae to many compounds have not yet been reported. One intriguing study discovered that bumblebees will avoid pollen that has been laced with the bitter compound, quinine (Muth et al., 2016a). Whether or not bumblebees can detect bitter compounds at their periphery and learn to associate peripheral taste stimuli, particularly that encountered when visiting flowers, with the presence or absence of reward has not yet been studied.

Here, we performed a series of experiments to explore the capacity of bumblebees to use peripheral taste to detect compounds on floral surfaces and use these to inform foraging choices. We tested whether freely flying bees have instinctual (spontaneous) preferences or aversions to compounds applied to the surfaces of artificial flowers. We also performed a free-flight learning assay to test whether bumblebees learn to use surface taste cues to predict the presence or absence of rewards provided by artificial flowers. We used sweet (sucrose), salty (NaCl) and bitter (quinine and caffeine) taste cues applied to the surfaces of artificial flowers in all experiments. We also used electrophysiological recordings of GRNs to test how bumblebees perceived these tastants in a subset of gustatory sensilla on the antennae. Such work expands our understanding of how peripheral taste can be is used by insects, and what kinds of chemical stimuli bees can detect via their peripheral taste systems.

## Materials and Methods

### Bumblebees

Buff-tailed bumblebees, *Bombus terrestris* subsp. *audax* (Harris 1776), were obtained from Biobest (Westerlo, Belgium, via Agralan, Swindon, UK). No ethical permissions were required for the experiments involving bumblebees, but the experiments were conducted according to ASAB/ABS guidelines. For electrophysiology and scanning electron microscopy, bumblebees were housed in their original colonies (as received) at RT (∼22°C) and provided with BioGluc sugar syrup (Biobest, Westerlo, Belgium) *ad libitum* , and honeybee-collected pollen (Agralan, Ashton Keynes, UK) three times per week. Only workers with thorax widths >4.5 mm were selected for experimentation to reduce the likelihood of nurse bee inclusion (Goulson et al., 2002).

### Scanning electron microscopy

We performed scanning electron microscopy (SEM) of the antennae of female worker bumblebees collected as exiting the colony. Bumblebees were cold-anesthetized prior to removing their antennae. Antennae were submerged in acetone for 24 h and then mounted on insect pins using UV-hardening dental glue. We sputter-coated the antennae with gold-palladium for 150 s at 18 mA and imaged with a Neoscope 2000 (Nikon Instruments, UK) at 10 kV high vacuum.

### Antennal sensillum recordings

Female worker bumblebees were collected as they exited the colony and anesthetized on ice. Each bee was secured in a copper tube to restrict movement, with dental wax stabilizing the head and preventing movement of the mouthparts. The antennae were oriented to expose the ventral side of the flagellum and affixed to the dental wax using wire staples. A silver wire reference electrode was inserted through a small puncture in the head capsule at the base of the eye and fixed with honeybee wax.

Electrophysiological recordings of gustatory sensilla were performed using tip recordings (Hodgson et al., 1955). A borosilicate glass recording electrode (15 μM tip diameter; Clark capillary glass GC150TF-10, pulled with a Narishige PC-10 electrode puller) containing the test solution was positioned over individual gustatory sensilla using a motorized micro-manipulator (MPC-200, Sutter Instruments, USA). Signals were acquired with a TasteProbe pre-amplifier (Syntech, Germany), amplified 100x using an AC amplifier (Model 1800, A-M Systems, USA; 100–5000 Hz bandpass filters), digitized at 30 kHz with a Data Translation DT9803 digitizer (Digilent, USA), and recorded in DataView v11.5 (St Andrews, UK).

Five test solutions (DI water, 100 mM sucrose, 100 mM NaCl, 1 mM quinine, and 1 mM caffeine) were presented in a pseudo-randomized order without the use of additional electrolytes (Kessler et al., 2015; Miriyala et al., 2018; Parkinson et al., 2022). Sensilla were stimulated with each solution for 3 seconds, with a minimum 3-minute inter-stimulus interval. Recordings were obtained from sensilla trichodea C/D located on the ventral surface of the distal flagellomere (mid and tip regions; Ågren, L. and Hallberg, E., 1996; Ren et al., 2023). For each bee, at least one mid and one tip sensillum was sampled, and in some cases, additional sensilla were recorded. A total of 26 sensilla (13 mid, 13 tip) were recorded across 10 individual bees. Sensilla that did not respond to all four of the tastants were not used in the analysis.

### Spike detection & analyses

Spike detection was performed in MATLAB (v R2020b, The MathWorks, MA, USA) pipeline to isolate and quantify neuronal activity while minimizing noise and artifacts. Spikes were detected by filtering the raw electrophysiological trace using a bandpass filter (100–1000 Hz) to isolate neural activity and normalizing the signal to remove baseline fluctuations. Peaks were identified using a peak-detection algorithm that located manually set threshold-crossing events corresponding to spike times. Spikes occurring within the desired time window (from 0.1 to 2.1 s after recording onset) were retained, and their waveforms (±2 ms around the peak) were extracted for further analysis. Spike artifacts were removed from the data in two steps. Spikes with peak amplitudes significantly higher than the mean were identified and excluded to eliminate large amplitude artifacts. Next, spikes with waveform widths exceeding 1.5 ms (45 samples at a 30 kHz sampling rate) were removed, as these were considered non-spike artifacts. After artifact removal, the remaining waveforms and spike locations were visually inspected to ensure the refinement process preserved valid spikes. Average spiking rates over the first 1 s were used for subsequent analyses.

To analyse average spiking rates from antennal recordings across stimuli and locations, the firing rate data (spikes/1s) was modelled in R version 4.3.3 (R Core Team, 2024) using Generalized Linear Models (GLMs) with various distributions due to non-normality detected by the Shapiro-Wilk test. Models with Gamma, Inverse Gaussian, and Gaussian distributions were compared using Akaike Information Criterion (AIC), with the Gamma model showing the best fit. Mixed-effects models were also tested, but the inclusion of random effects for "Bee ID" or "sensillum ID" did not significantly improve model fit and were therefore excluded. The final Gamma GLM included interaction terms for stimulus and sensillum location (tip or mid). A Wald test was conducted to evaluate the overall effects of these factors using the ‘car’ package (Fox and Weisberg, 2011), followed by post hoc pairwise comparisons using estimated marginal means (emmeans) from the ‘emmeans’ package (Lenth, 2020).

We also compared the temporal GRN responses over 2 s by averaging spikes in 100 ms bins (20 bins in total). To analyse the effects of tastant type and time on the spiking responses, a generalized linear mixed-effects model was fitted to the data from the mid-location sensilla using the ‘glmmTMB’ package (Brooks et al., 2017). The model assumed a Gamma distribution with a log link function to account for the non-negative and potentially skewed distribution of spike counts (+1 was added to each bin to account for zeros). The fixed effects included stim, time, and their interaction, while the “bee ID” was included as a random effect to account for repeated measures from the same individuals. Model fit was assessed using a type II ANOVA to evaluate the significance of the fixed effects. Post hoc pairwise comparisons were conducted with emmeans to identify differences between stimuli at each time point. This approach allowed for a robust examination of temporal and stimulus-specific variations in spiking activity between tastants.

### Free flying bumblebee conditions

Upon arrival, bumblebee colonies were re-housed in multi-chambered wooden colony boxes with transparent Perspex lids. Each box contained a ‘colony chamber’ where the colony was placed, and a ‘connecting chamber’. Colony boxes were lined with baking paper and kitty litter (CATSAN hygiene+, CATSAN, Cork, Ireland). Colonies were provided with cotton wool bedding.

Bees accessed a flight arena (70 x 100 x 30 cm, width x length x height) via a clear access with gated controls. The arena had a clear, UV transmissive Perspex lid, a green painted floor (PlastiKote Fast Dry Project Enamel “Garden green, B9”, Motip Dupli B.V., Wolvega, The Neatherlands), and six experimenter access doors. Arena and colony connecting chambers were illuminated by daylight bulbs on an automated 08:00-20:00 h day-night cycle.

Bee colonies were fed honeybee-collected pollen (Agralan) three times per week and had *ad libitum* access to 1M sucrose solution provided in artificial flowers and PCR racks placed haphazardly within the arena. These artificial flowers, similar in design to experimental ones, consisted of specimen jars (white screw cap 60 ml PP Container, Greiner bio-one, Stonehouse, UK) with upturned 1.5 ml centrifuge tube lids stuck to the lid as feeding wells.

Foraging adult workers, identified while feeding from artificial flowers, were marked with queen marking plates (Lyson Beekeeping, Klecza Dolna, Poland). Before experimentation, the flight arena was cleared, feeders removed, and the arena floor cleaned with 50% ethanol solution and dried. Bees were restricted to the colony and connecting chambers until experiments began.

### Artificial flower design

Artificial flowers with controlled surface chemistry were used in behavioural experiments (Fig. 1). Each flower consisted of a 60 ml specimen jar (white screw cap 60 ml PP Container, Greiner bio-one, Stonehouse, UK) with its screw cap serving as the flower top. A 0.2 ml centrifuge tube lid, cut free and upturned, served as a feeding well containing sucrose solution or water. The jar body was wrapped in black electrical tape (2Brothers, Preston UK) to prevent visual distractions.

**Figure 1:**
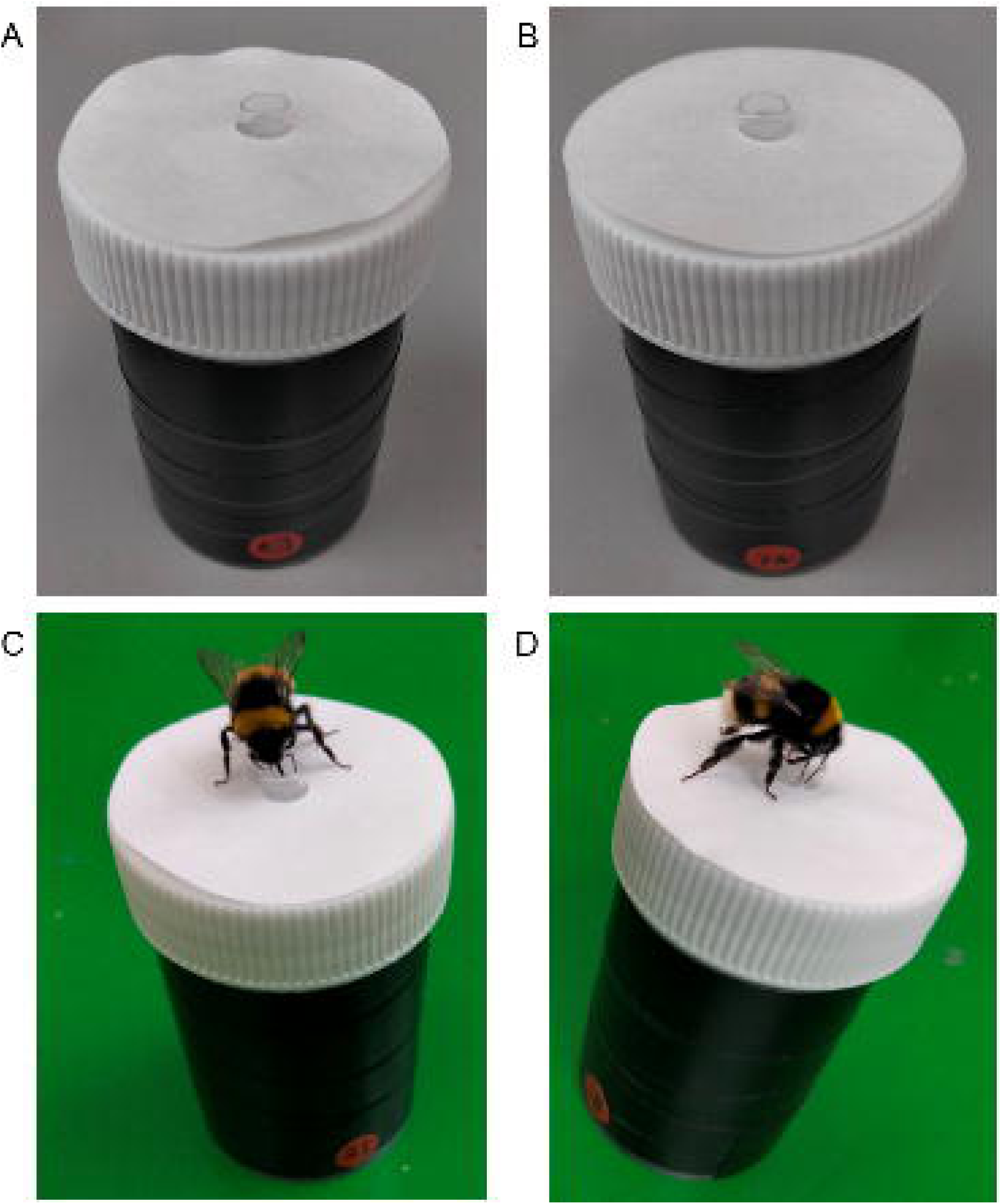
Examples of the artificial flowers used in bee behavioural experiments. A) An untreated flower, B) a Sucrose treated flower. C) and D) bumblebees feeding from artificial flowers. Note the numbered markings on the flower bases.

A 42.5 mm filter paper disc (VWR international, Radnor, Pennsylvania, USA) was attached to the flower top using double-sided tape (3M Scotch, Radnor, Pennsylvania, USA). A 6 mm hole punched in the centre of the disc allowed it to fit around the feeding well. Discs were treated with a tastant (sucrose, sodium chloride [NaCl hereafter], quinine hydrochloride [quinine hereafter], or caffeine) or left untreated. All flower variants were visually identical. To aid identification of flowers by experimenters without providing an additional cue to the bees, red sticky dots were stuck about the base of the flower, and two-digit numbers written on these with black permanent marker (Fig. 1).

### Flower disc preparation

Discs were prepared in three steps: chemical application, drying, and final assembly. Batches of 300- 400 discs were processed at a time. If discs treated with a surface tastant were prepared, similar quantities of untreated discs were also prepared at the same time. Discs of separate treatments and untreated discs were kept separate throughout preparation (separate storage containers, foil sheets etc., see below) even if prepared at the same time. All discs were handled using nitrile gloves, which were changed regularly and between different treatments, and tweezers, which were cleaned regularly and between treatments with RO Water. Irregular, creased or damaged discs were not used. Likewise, discs that, prior to application to artificial flowers, were discarded if they came into contact with surfaces other than those detailed below or clean paper towel laid out on surfaces. These procedures avoided introduction of specific differences or contaminants to individual discs, and the flowers they are attached to, which may be utilised by bees to inform foraging.

To apply tastants, filter paper discs were submerged in a petri dish containing the treatment solution until saturated (∼0.3 ml per disc). Solutions (0.5 M for sucrose and NaCl; 1 mM for quinine and caffeine) were freshly prepared using RO water (<50 µS, Elga Purelab Chorus 3, Veolia Water, Paris, France). Untreated disks were submerged in pure RO water but otherwise prepared in the same way.

Treated discs were dried in a climate chamber (Memmerts HPP750, Schwabach, Germany) on tin foil at 55 °C and 20 % humidity for 45-90 minutes until fully dry. Once dried, discs were stored in sealed plastic boxes by treatment type. Dry discs had a hole punched in their centre using a 6 mm single point hole punch (G4Gadget, Thurmaston, UK). Then to one side of the disc, two sections of double-sided tape of approximately 1 cm^2^ would be stuck either side of this hole. Discs were then temporarily stuck down into the inside of A4 booklets of tracing paper that would be stored in sealed plastic containers to await experiments. Prior to presentation of flowers to bees, the discs were stuck directly onto artificial flower tops (Fig. 1). Due to the finish of the tracing paper, discs and their attached tape could be removed cleanly with minimal residue.

### Controlling for UV and surface texture differences

Although all artificial flower variants appeared uniformly white, bees can detect UV reflectance (Davies et al., 2013; Dyer and Chittka, 2004b). Spectral analyses showed small UV reflectance differences between untreated and sucrose, caffeine, and quinine-treated flowers, but not NaCl-treated flowers (supplementary material 1A). However, these differences were minor and unlikely to affect foraging behaviour (see Dyer and Chittka, 2004b).

As bees can detect surface texture cues, this was examined using SEM (Supplemental material 1B). No meaningful structural changes were observed for sucrose, quinine or caffeine treatments, with all discs showing the natural random fibre structure of filter paper. NaCl-treated discs exhibited small, irregular residues (Fig. S2), but these were below the size (<5 µm) typically detectable by bees (Kevan and Lane, 1985; Whitney et al., 2009; Whitney et al., 2011). Based on SEM imaging, chemical treatments did not alter the surface texture in a way that bees could detect (Fig. S2).

### Artificial flower cleaning during experiments

Artificial flowers were cleaned with 50% ethanol before disc attachment and at the end of trials. In the differential conditioning experiments, where a ‘fresh set’ of artificial flowers were presented in a test phase (see below), this separate fresh set were also cleaned before disc attachment and presentation to bees.

During experiments, flowers were cleaned after each foraging bout (the time between a bee’s departure and return to the nest), unless the bee made fewer than 6 flower visits in the bout (visits defined below) and the flowers had already been cleaned beforehand. This minimized scent marks accumulation, which could influence foraging choices (Pearce et al., 2017; Stout and Goulson, 2001), while reducing disc waste.

Cleaning involved removing flowers from the arena, wiping feeding wells with paper towel, discarding used discs, unscrewing plastic lids, and washing them with 50% ethanol before drying. Lids were then randomly reassigned to flower bases. Discs from the same preparation batch were distributed equally among flower variants to prevent contamination bias. If flowers did not require full cleaning, feeding wells were still simply emptied and refilled to prevent evaporation differences affecting foraging choices. Any visibly soiled discs (e.g., due to defecation or dirt) were replaced.

### Bee behavioural experiments

Two experiments were conducted: preference experiments (e.g. Dyer et al., 2006; Lehrer et al., 1995; Raine and Chittka, 2007) and differential conditioning (learning) experiments (e.g. Dyer and Chittka, 2004a; Kunze and Gumbert, 2001; Raine and Chittka, 2008).

Trials were not time-limited but measured by the number of flower visits, as this better reflects exposure to artificial flower cues (e.g. Clarke et al., 2013; Dyer and Chittka, 2004a; Raine and Chittka, 2007; 2008). However, all trials took place between 09:00 and 20:00 h, and individual bees completed their trial on the same day they started. Bees that failed to complete the experiment were not reused, and their data were excluded.

If a bee became visibly injured during a trial, the trial was terminated and its data excluded. This occurred only once in the conditioning experiments. Each bee participated in only one the experiment, and one treatment group to maintain independent observations.

### Surface tastant preferences

In spontaneous preference experiments, bees were presented with 12 artificial flowers, each providing 25 µl of 1 M sucrose solution in the feeding well. Each bee was assigned to one of four tastant treatments (sucrose, NaCl, quinine, caffeine), determining the surface chemistry of its flowers presented to the bee. Half of the (12) flowers were treated with the corresponding tastant, while the other half were untreated, arranged randomly in the arena with at least 10 cm spacing. Sucrose solutions were stored at 4 °C for up to one week. Lab conditions were maintained at 23.23°C ± 0.17°C and 47.75% ± 0.69% humidity (mean ± s.e.m.).

Marked bees were tested individually in the arena, free to forage on the artificial flowers. Typically, upon encountering artificial flowers, a foraging bee slowed down flight and made contact with the tops of artificial flowers with its feet for landing. After landing, a bee either: a) fed by extending its proboscis into the feeding well (‘feeding’, as in Fig. 1C-D); or b) departed without feeding. We monitored which flowers were visited and whether the bee fed during the flower visit or departed without feeding. A bee was considered to have landed on flower if the bee touched the top of the flower with its feet. If the bee made contact with a flower surface with it’s feed (as in the above description) but did not completely quit flight before departing, such a visit was still counted as a landing and departure without feeding. This criteria for classification of when a landing and flower visit as such has been used previously in comparable studies (Harrap et al., 2017; 2019; 2021). We also noted whether bees contacted the flower disc with their mouthparts, to evaluate instances where the bee could use oral taste to inform decisions.

After each flower visit, while the bee was elsewhere in the arena, the flower was removed, refilled, and repositioned (>10 cm from other flowers). If a bee revisited a flower before relocation, the visit was not counted. Bees could return to the nest at any time and were re-released for another foraging bout. Between bouts, all flowers were cleaned, discs replaced if needed, and repositioned.

The experiment ended when the bee made 20 flower landings. A total of 48 bees (12 per tastant) from three colonies were tested.

For each flower visit made we determined whether the bee demonstrated a response in favour of treated flowers or not (as in Dyer et al., 2006; Harrap et al., 2021; Lawson et al., 2018; Raine and Chittka, 2007):

- Flower visits in favour of treated flowers: visiting a treated flower and extending the proboscis into the feeding well (positive selection of those flowers), or visiting an untreated flower and leaving without extending the proboscis into the feeding well (a rejection of flowers lacking that surface tastant).
- Flower visits not in favour of treated flowers: visiting an untreated flower and extending the proboscis into the feeding well, or visiting a treated flower and leaving without extending the proboscis into the feeding well.

Bees may have landed on flowers for other reasons than seeking rewards, such as grooming, that may result in rejections not indicatory of foraging preferences toward flower surface chemistry. However, such flower landings are rare and would be as likely to occur on either flower variant.

To quantify preferences, we calculated a Tastant Surface Response Rate, the proportion of landings favouring treated flowers. If foraging was random, this rate would be 0.5. Since response rates were bounded between 0 and 1, they were arcsine square-root transformed for statistical analysis. For each tastant, a two-tailed one-sample Wilcoxon signed-rank test was performed in R (version 4.1.3, R Core Team, 2022) to compare the median response rate against the expected random choice value (0.79 after transformation).

### Differential conditioning of surface tastants

The differential conditioning experiment consisted of two phases: training and test. Lab conditions were maintained at (mean ± s.e.m.) 22.56°C ± 0.15°C and 41.52% ± 0.38% humidity.

### Training phase

In the training phase, individually marked forager bees were tested alone in an arena with 12 artificial flowers placed randomly (>10 cm apart): Six rewarding flowers containing 25 µl of 1M sucrose solution (the unconditioned stimulus); and six nonrewarding flowers containing 25 µl of water (potentially an aversive ‘punishment’). Flowers were visually distinguished using even-numbered (rewarding) and odd-numbered (non-rewarding) labels. Solutions used were stored at 4°C for up to one week.

Bees were assigned to one of nine conditioning groups (Table 1). This determined the surface chemistry of the rewarding and nonrewarding flowers presented to bee. These included a ‘tastant- positive’ (rewarding flowers had the surface tastant, non-rewarding flowers were untreated) and ‘tastant-negative group’ (non-rewarding flowers had the tastant, rewarding flowers were untreated) for each tastant, and control group (both flower types were untreated). Each group had 10 bees (90 total) from seven colonies, with two colonies also used in preference experiments.

**Table 1:** Summary of the nine conditioning groups used in the differential condition experiments.

| Conditioning group | Type of discs applied to... |  |
| --- | --- | --- |
|  | Rewarding flowers | Nonrewarding flowers |
| Control | Untreated | Untreated |
| Sucrose positive | Sucrose | Untreated |
| Sucrose negative | Untreated | Sucrose |
| NaCl positive | NaCl | Untreated |
| NaCl negative | Untreated | NaCl |
| Quinine positive | Quinine | Untreated |
| Quinine negative | Untreated | Quinine |
| Caffeine positive | Caffeine | Untreated |
| Caffeine negative | Untreated | Caffeine |

Individual bees were allowed to forage freely on these artificial flowers. We monitored both whether the bee visited rewarding or non-rewarding artificial flowers (visit as defined in the preference experiment) and whether the bee extended its proboscis into the feeding well (‘drinking’), or left without doing so at each flower visit. Additionally, at each visit we monitored whether the bee brought its mouthparts into contact with the flower disc surface. As the bee foraged, flowers were moved and refilled as described for the preference experiment.

To minimize scent mark accumulation, flowers were removed after a bee’s twentieth visit in a single foraging bout. This would be done carefully while the bee was elsewhere in the arena. Consequently, it was not always possible to achieve immediately following the 20^th^ visit of a bout (max bout length achieved: 28 visits). With flowers removed bees either returned to the nest voluntarily or were manually guided back. The training phase continued each bee completed 70 flower visits, sufficient for learning other floral cues (see, Clarke et al., 2013; Dyer and Chittka, 2004a; Harrap et al., 2017; 2019; 2021; Whitney et al., 2008).

### Test phase

After completing 70 visits, bees entered the test phase on their next foraging bout. Here, a fresh set of artificial flowers, identical to those in training, was presented to the bee. However here but all contained only 25 µl of water in their feeding wells. Bees were allowed to forage freely for 20 visits, and behavior was monitored as in training, where whether a flower variant had been rewarding and nonrewarding in the training phase was used to determine a flower’s reward state. For the control group, flowers were distinguished only by their numerical labels, consistent with their previous reward status. Flowers were moved, refilled and repositioned as in the training phase.

### Data analysis

For each flower visit in both phases, decisions were classified as correct or incorrect (as in Dyer and Chittka, 2004a; Harrap et al., 2021; Raine and Chittka, 2008):

- Correct: Landing on a reward flower and extending the proboscis into the feeding well, or landing on a non-rewarding flower and leaving without extending the proboscis into the feeding well
- Incorrect: Landing on a non-rewarding flower and extending the proboscis into the feeding well, or landing on a rewarding flower and leaving without extending the proboscis.

Non-foraging landings (e.g., grooming) were rare and unlikely to bias results (see above).

The ‘success rate’ (proportion of correct decisions) was calculated at 10-visit intervals (e.g., after 10, 20, 30 visits) during training and across the 20 visits of the test phase. This metric follows previous work (see, Dyer and Chittka, 2004a; Harrap et al., 2021; Raine and Chittka, 2007). A bee randomly foraging bee would have a success rate of 0.5.

Success rates of the control group were compared to the tastant-positive and tastant-negative groups for each tastant. Arcsine square-root transformation was applied to account for data bounded between 0 and 1. Generalised linear models (GLMs) were fit to data using R version 4.3.2 (R Core Team, 2022), with AIC model selection (Richards, 2008) identifying the most parsimonious model. The modelling approach followed Harrap et. al. (2017; 2020; 2021), and is described in Supplementary Materials 1C. For the test phase, success rates were compared across tastant- positive, tastant-negative, and control groups of each tastant using ANOVA, with *post hoc* Tukey tests, in R.

## Results

### Bumblebee antennal gustatory sensilla respond to bitter compounds

Scanning electron microscopy (SEM) identified trichodea C/D sensilla on the ventral mid-surface and the tip of the distal antennal flagellum (Fig. 2A-C), known to house four gustatory receptor neurons (GRNs; Ågren, L. and Hallberg, E., 1996). Using tip recordings (Hodgson et al., 1955), we tested responses to DI water, 100 mM sucrose, 100 mM NaCl, 1 mM quinine, and 1 mM caffeine, observing spiking activity for all stimuli (Fig. 2D).

**Figure 2:**
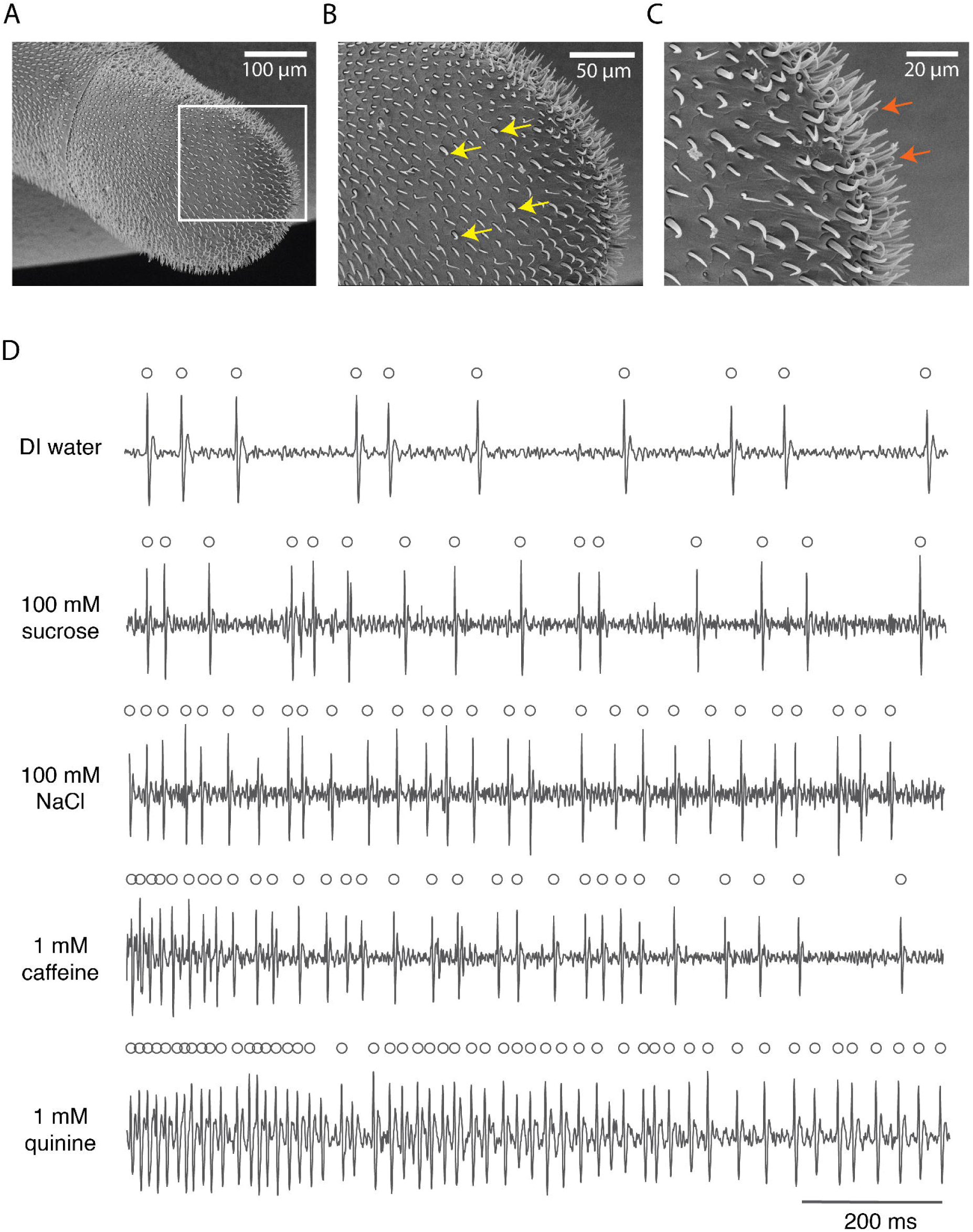
SEM images and electrophysiological responses of gustatory sensilla on the bumblebee antenna. A-C) SEM images of the ventral surface of the distal flagellum revealed trichodea C/D sensilla on the mid (B) and tip (C) of the flagellomere. D) Representative electrophysiological traces from a tip sensillum showing GRN spiking when stimulated with DI water, 100 mM sucrose, 100 mM NaCl, 1 mM caffeine, or 1 mM quinine.

The sensilla on the antennae that we recorded from housed at least one sugar-sensing GRN, one salt-sensing GRN, one water sensing GRN, and one bitter sensing GRN (Fig 2D). The bitter-sensing GRN in each sensillum spiked in response to stimulation with quinine or caffeine (Figure 3). GRN spiking rates varied significantly by stimulus type, recording location, and their interaction (stimulus: χ² = 189.6, p < 0.001; location: χ² = 9.44, p < 0.001; interaction: χ² = 22.3, p < 0.001, Fig. 3A). Water elicited the lowest spiking responses. The mid sensilla responded with higher average firing rates to all other stimuli compared with water, although between the other tastants there were no significant differences in average firing rates. Notably, the tip sensilla exhibited the highest spiking rates for caffeine and quinine, suggesting that the bitter-sensing GRNs in this location were particularly sensitive to these alkaloids.

**Figure 3:**
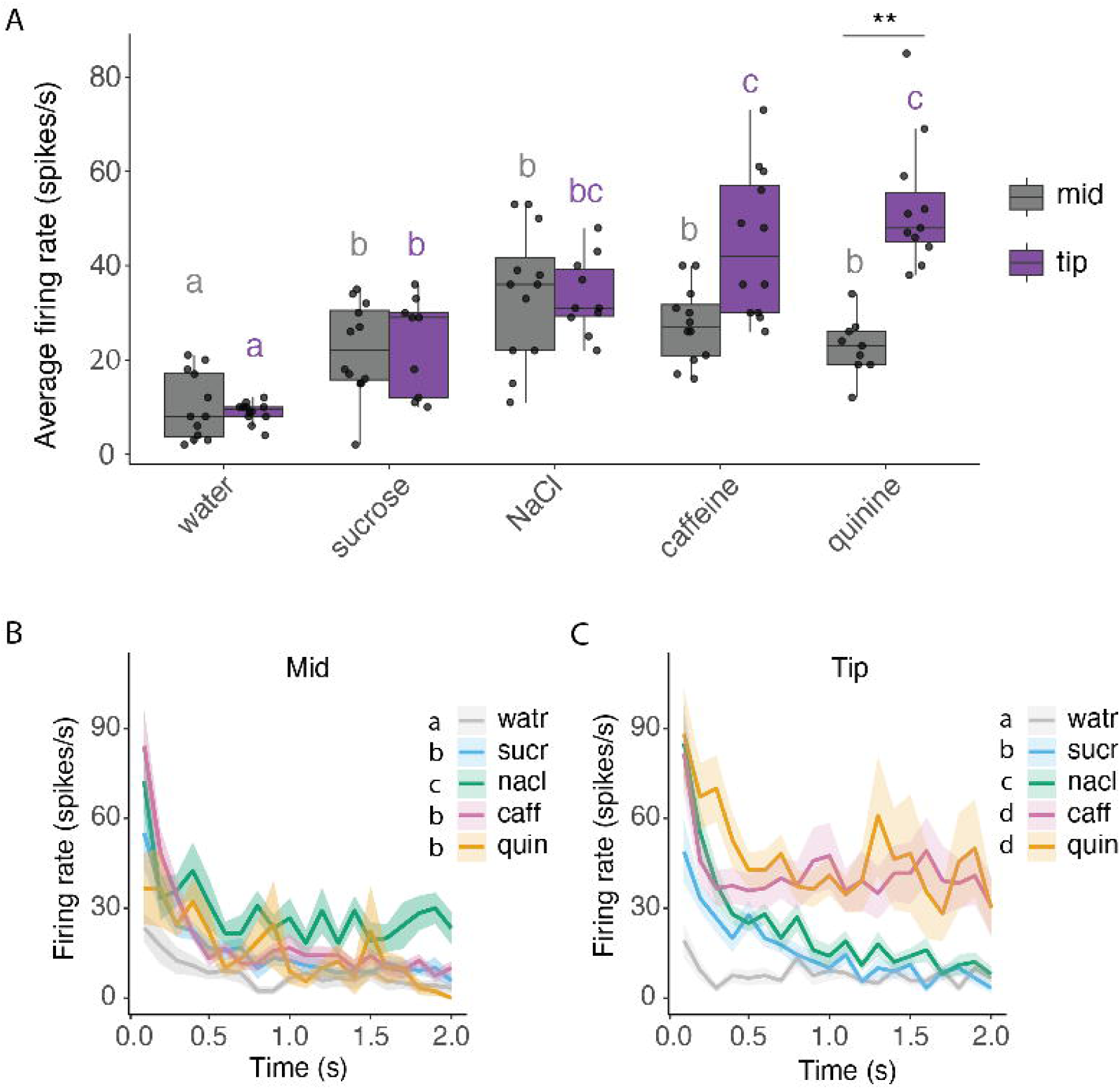
Antennal GRN responses to tastants vary by sensillum location. A) There were significant differences in the GRN firing rates over 1 s of stimulation (n = 11-13 sensilla per stimulus). Boxplots show median, 25^th^ and 75^th^ percentiles, with whiskers showing the range. Filled circles show the responses of individual sensilla. Results from an emmeans *post hoc* test noted, with significant effects across stimuli within each location denoted with coloured letters, and differences between locations for a given stimulus shown with asterisks. B-C) Responses of GRNs from sensilla in the mid (B) and tip (C) locations over 2s of stimulation (n = 11-13 sensilla per stimulus). Lines show mean firing rates across all replicates with standard error shading. Results from emmeans *post hoc* test displayed as letters beside legend.

The tastant used to stimulate the GRNs influenced the temporal pattern of spiking as a function of time at the tip and on the ventral mid-surface of the antennae (LME, mid location: stimulus χ² = 182.1, p < 0.0001; time χ² = 190.8, p < 0.0001; interaction χ² = 21.2, p = 0.0003; tip location: stimulus χ² = 497.5, p < 0.0001; time χ² = 128.6, p < 0.0001; interaction χ² = 53.3, p < 0.0001). GRN spiking responses displayed phasic-tonic activity, with initial high firing rates declining over 0.4 s before stabilizing. Caffeine and quinine sustained significantly higher tonic firing rates over 2 s at the tip (emmeans, p < 0.0001), reinforcing their role in bitter detection (Fig. 3 B-C).

### Naïve bumblebees show no floral preferences based on surface chemistry differences

In preference experiments, we tested whether naïve bumblebees preferred or avoided flowers with additional surface tastants (sucrose, NaCl, quinine or caffeine) compared to untreated flowers. Of 86 bees tested, 70% (60/86) began to visit the flowers, 80% of these (48/60) completed 20 visits and were included in the analysis. Bees completed the preference experiment in 57 ± 4 mins (mean ± s.e.m.) across 3.29 ± 0.16 foraging bouts with 6.08 ± 0.29 visits per bout. Few bees contacted the flower discs with their mouthparts (7 bees total), with one bee performing two such visits, the other six performing only one such visit.

Naïve foragers showed no significant preferences for treated or untreated flowers (Fig. 4), with equal visit frequencies across flower types, regardless of the surface tastant. Tastant surface response rates did not differ from random choice (0.5, Table 2).

**Figure 4:**
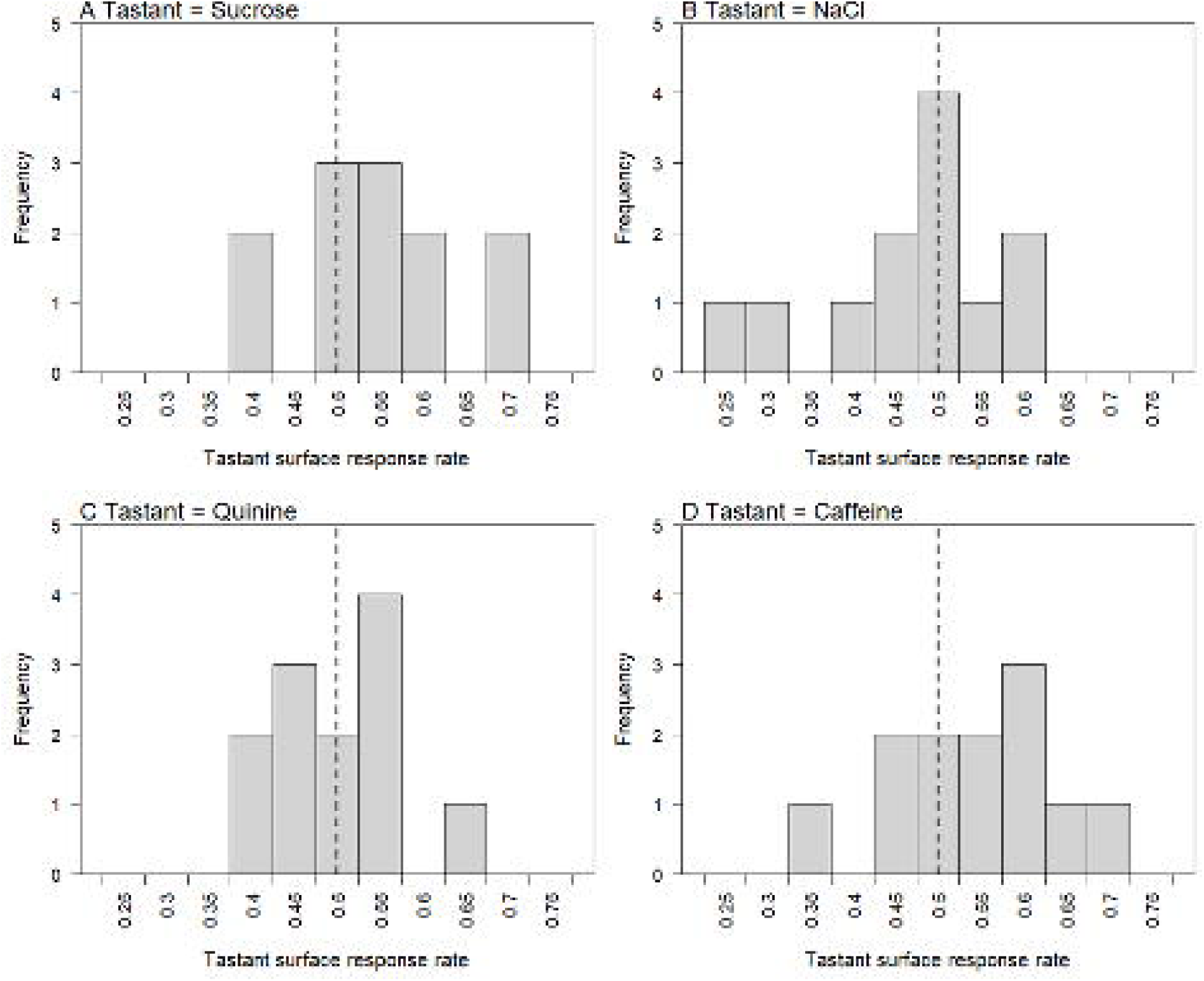
Histogram showing bee responses to artificial flowers in the preference experiments. Bars represent frequency of bees (*n*=12 in each assigned tastant) that over 20 flower visits achieved each tastant response rate, indicating the proportion of visits where responses showed favour of flowers with surface chemistry treated with A) Sucrose, B) NaCl, C) Quinine and D) Caffeine, over control flowers. Dashed vertical line indicates the expected tastant surface response rate of bees that forage randomly (0.5).

**Table 2:** Summary of the Wilcoxon signed rank tests performed on bee responses in preference experiments. ‘Tastant’, corresponds to that applied to treated flowers presented to bees, *n*=12 bees for each tastant.

| Tastant | Tastant surface response rate |  |  | *W* | *P* |
| --- | --- | --- | --- | --- | --- |
|  | Median | Minimum | Maximum |  |  |
| Sucrose | 0.55 | 0.40 | 0.70 | 53 | 0.288 |
| NaCl | 0.50 | 0.25 | 0.60 | 22 | 0.194 |
| Quinine | 0.50 | 0.40 | 0.65 | 30 | 0.502 |
| Caffeine | 0.55 | 0.35 | 0.70 | 53 | 0.288 |

### Bumblebees learn to favour rewarding flowers based on surface chemistry

In the differential conditioning experiment, we tested whether bees could learn to associate surface tastants with reward. Of 176 bees tested, 75% (132/176) visited flowers, and 68% of these bees (90/132) completed the experiment. Bees completed 70 training visits in 5.91 ± 0.14 foraging bouts (mean ± s.e.m.), with 11.84 ± 0.21 landings per bout. All but one bee completed the test phase in a single bout. However, this bee which required two bouts made only 5 visits in the first of these. Consequentially, all bees completed the test phase on a single ‘set’ of flowers (i.e. without cleaning). The experiment lasted 2 h, 24 mins ± 4 mins per bee. Mouthpart contact with flower discs was rate (seen in 14 visits by 12 bees).

Control bees (i.e. those tested with flowers identical in surface chemistry) learned to favour rewarding flowers in training, increasing success rate from 0.45 to 0.57 after 70 visits (Fig. 5, black lines). However, in the test phase, when presented with a fresh set of flowers, their success dropped to 0.49, showing frequencies of correct visits comparable to random foraging (Wilcoxon signed rank test, conducted as in preference experiments on control group test phase success rates, *W*=15, *n*=10, *P*=0.22).

**Figure 5:**
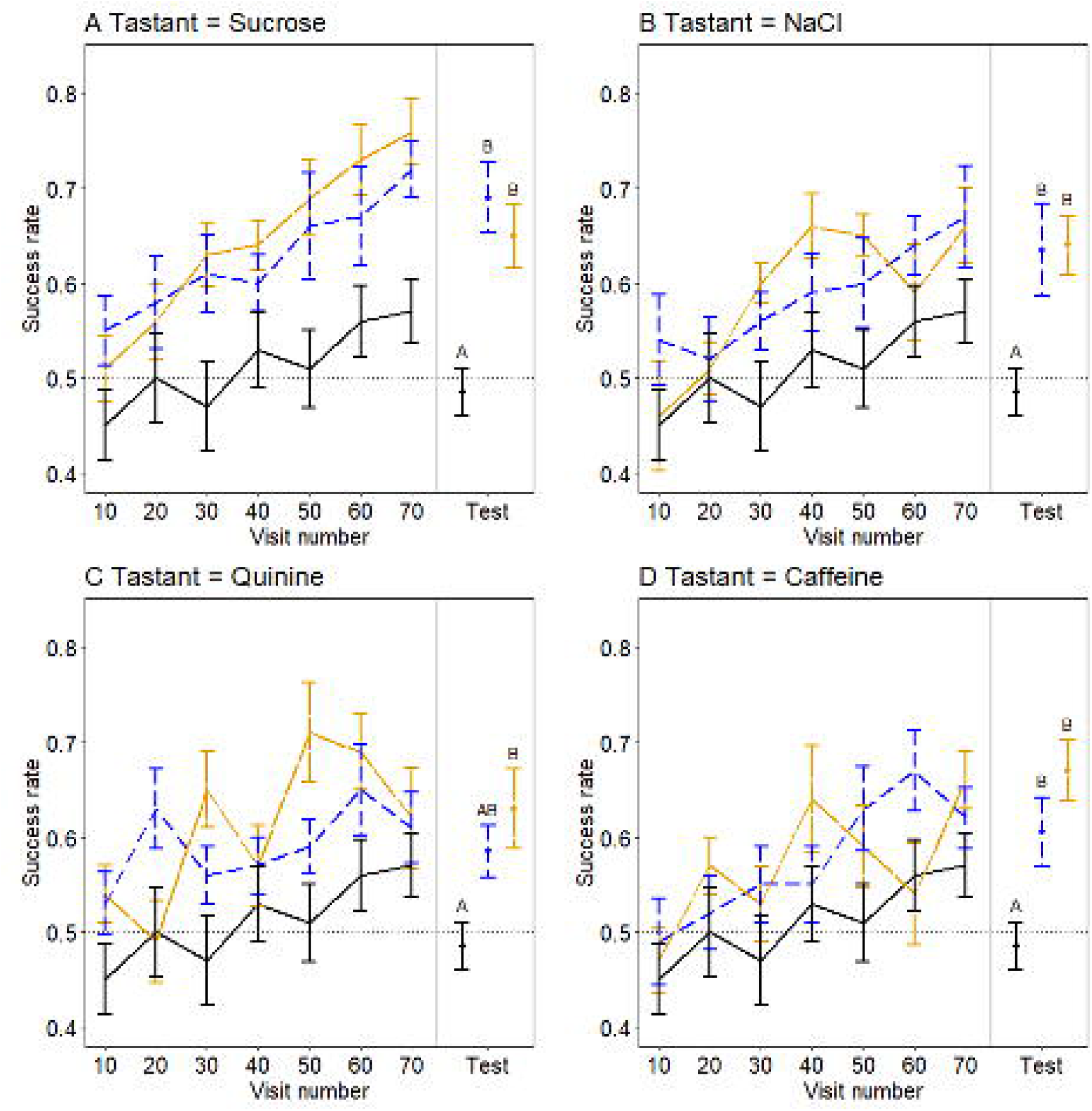
The relationship of bees foraging success and experience of artificial flowers differing in A) Sucrose, B) NaCl, C) Quinine and D) Caffeine surface chemistry cues. Solid lines indicate mean success rate (proportion of ‘correct’ visits with regards to floral rewards) of bees in the previous 10 visits, calculated at 10 visit intervals. Also plotted are the mean success rates of bees in each conditioning group, during the test phase, mean value indicated by points above ‘Test’. Error bars represent ±s.e.m throughout. Colour and dashing of lines indicate conditioning groups of each tastant. Solid black line indicates the control group, where no difference in surface chemistry is present (this conditioning group is copied across panels to facilitate comparisons). Blue dashed line, the tastant positive group, where rewarding flowers are treated with the surface tastant. Orange longdashed line, the tastant negative group, where nonrewarding flowers are treated with the surface tastant. Letters above mean success in the test phase represent grouping according to Tukey’s *post hoc* tests, groups that share letters show no significant difference (p<0.05) from groups with the same letter. Values of *n* for each conditioning group of each tastant treatment is 10 bees. Dashed horizontal line indicates the expected success rate of bees that forage randomly (0.5).

### Bees remember a sucrose or a salty (NaCl) surface taste cue associated with reward presence and absence

During the training phase bees in the sucrose groups learned to favour rewarding flowers, reaching >0.7 success rate by the end of training (Table S2, Fig. 5A). In testing, sucrose- conditioned bees still preferred flowers matching their training rewards, outperforming controls (ANOVA, *F*_2,27_=10.91 *P*<0.001). There was no difference between test phase success rates of sucrose-positive and sucrose-negative groups (Tukey’s *post hoc*: *P*=0.63), but both differed from controls (versus sucrose-positive: *P*<0.001; versus sucrose-negative: *P*<0.01).

During training NaCl-negative bees (non-rewarding flowers treated) outperformed controls, while NaCl-positive bees (rewarding flowers treated) performed similarly to controls (Table S3, Fig. 5B). Bees in both NaCl conditioning groups achieved success rates of ∼0.6 after 70 flower visits. Both NaCl groups showed learned preferences in the test phase, achieving higher success than controls (ANOVA, *F*_2,27_=5.64 *P*=0.01; Tukey’s *post hoc* : control vs. NaCl positive *P*=0.02; control vs. NaCl negative *P*=0.02; NaCl positive vs. NaCl negative *P*=1.00).

### Bees find it difficult to associate quinine with reward but not caffeine

Quinine-positive bees (rewarding flowers treated) outperformed controls in training, while quinine- negative bees (non-rewarding flowers treated) did not (Table S4, Fig. 5C). Bees in both quinine conditioning groups achieved success rates of ∼0.6 by the end of training. However, in testing, quinine-negative bees successfully avoided quinine flowers, outperforming controls (ANOVA, *F*_2,27_=5.20 *P*=0.01; Tukey’s *post hoc*: control vs quinine negative, *P*=0.01). Quinine-positive bees failed to favour quinine-treated rewarding flowers, showing performance comparable to both control and quinine-negative groups (Tukey’s *post* hoc: control vs. quinine positive *P*=0.11; quinine positive vs. quinine negative *P*=0.55).

Bees in caffeine groups performed like controls in training but reached >0.6 success rate after 70 visits (Table S5, Fig. 5D). In testing, caffeine-conditioned bees still favoured previously rewarding flowers, achieving higher success than controls (ANOVA, *F*_2,27_=8.76 *P*=0.001). There was no difference between caffeine-positive and caffeine-negative groups (Tukey’s *post hoc* : caffeine positive and caffeine negative *P=*0.32), but both differed from controls (Tukey’s *post hoc*: control vs. caffeine positive *P*=0.03; control vs. caffeine negative *P*<0.001).

## Discussion

Our data are the first to show that bumblebees can learn to associate compounds detected on the surfaces of flowers using their peripheral taste system (antennae and/or tarsi) with the presence or absence of rewards. Throughout these experiments, the bees rarely brough their proboscis into contact with flower surfaces, indicating that oral mechanisms for tasting flower surface chemistry and distinguishing flowers was not used. Naïve bees showed no instinctual responses toward or against flowers with any surface cue we tested. However, in the learning experiments, bees could learn and remember the presence or absence of rewards when it was associated with sucrose, NaCl, or caffeine surface chemistry. While they could learn to associate quinine surface chemistry with the absence of rewards, it was harder for them to remember that it was associated with rewards during the test phase. Thus, we are the first to show that like mammals (e.g. Appleton et al., 2018; Capaldi et al., 1997; Garcia-Burgos and Zamora, 2015), bees can overcome instinctual aversions to tastes and learn to associate them with rewards. The difficulty associating bitter surface chemistry with rewards could relate to a compound’s toxicity (Ayestaran et al., 2010; Barlow et al., 2017; Tiedeken et al., 2014; Villalona et al., 2020; White and Chapman, 1990). Intriguingly, bumblebees could also associate the taste of nutrients with the absence of rewards.

Bumblebee floral surface chemistry learning was most clear within the test phase of our conditioning experiment, but less pronounced during the training phase. Several tastant conditioning groups that showed enhanced foraging success in the test phase showed comparable success to the control group during the training phase (see tables S2-S5). The difficulty of the given learning task may explain this inconsistency, as the sucrose conditioning groups showed enhanced foraging success throughout the experiment. However, it is most likely the result of learning occurring in control group bees over the training phase, evidenced by their success rates >0.5. Control group bees may have learnt subtle differences between individual flowers (discussed in Harrap et al., 2021), or more likely scent marks left by the bees (Stout and Goulson, 2001). Filter paper has previously been used to collect bee scent marks and retains them well (Pearce et al., 2017). While discs were replaced regularly (see methods), deposition of scent marks between cleaning might still allow bees to temporary identify rewarding flowers once refilled and returned to the arena. When surface chemistry differences are present, bees may not attend to scent marks, or the presence of both cues does not result in a detectable improvement in foraging decisions (as in Harrap et al., 2019; Lawson et al., 2017). Hence, bees may not always show higher success rates than the control group during training despite forming associations between flower surface chemistry and rewards. In the test phase, the fresh set of artificial flowers lack the same individual identifiers and that scent marks. Regardless, the control group lacked whatever cues they had been utilizing in training, and reverted to random foraging. Meanwhile, conditioning group bees still had access to surface chemistry cues they had learnt. Thus, the difference in performance (when present) between control and conditioning groups, and thus floral surface chemistry learning, is more apparent in the test phase.

We show bumblebees have a robust capacity to learn based on peripheral taste stimuli using contact chemoreception. The role of peripheral sucrose stimulation in food choice has been demonstrated in restrained and walking honeybees and bumblebees (Mommaerts et al., 2013; Simcock et al., 2018 and others cited above). Honeybees have also been shown to be able to learn odours based solely on peripheral stimulation with sucrose or pollen (Bitterman et al., 1983; de Brito Sanchez et al., 2008; Nery et al., 2020; Wright et al., 2005a; Wright et al., 2005b). Restrained honeybees have similarly been observed to respond to peripheral stimulations with salt solutions (Bestea et al., 2021; de Brito Sanchez et al., 2014; Lau and Nieh, 2016). However, use of dry surface chemistry detected by peripheral taste as a cue, learnt separately from food or water consumed, as we show here, has been less frequently seen in bees. Honeybees have previously been shown to associate peripheral stimulation with sucrose and salt solution with an electric shock punishment, but not quinine solution (Guiraud et al., 2018). Bumblebees in our experiments were able to use the peripheral detection of sucrose and salt with the absence of reward. That the taste of a nutrient can become a learned cue of the absence of rewards could be similar to the type of learning mechanisms that underly conditioned food aversions reported in vertebrates (Nachman and Hartley, 1975; Parker, 2014; Wise et al., 1976).

Bees can evaluate and learn associations between floral cues and pollen rewards surface chemistry cues detected via peripheral taste have been proposed to have a role in this (Muth et al., 2015; Muth et al., 2016b; Nebauer et al., 2023; Nery et al., 2020; Nicholls and de Ibarra, 2014; Nicholls et al., 2019; Ruedenauer et al., 2015; Ruedenauer et al., 2017; Ruedenauer et al., 2019; Ruedenauer et al., 2021). Peripheral taste would be particularly useful in this scenario as foraging bees typically do not consume pollen (Heinrich, 2004; Nicholls and Hempel de Ibarra, 2017), but it would mean that bees do not need to associate post ingestive feedback about nutrients or toxins with the sensation of taste. Our findings confirm bumblebee peripheral taste capacity is sufficient to guide such foraging decisions, bees being able to associate surface chemistry distinct from consumed food with a positive or negative outcome. Furthermore, bumblebees have been reported to show aversion towards, spending less time collecting and grooming, alkaloid (Jacquemart et al., 2019; Muth et al., 2016a) and saporin (Wang et al., 2019) laced pollen. Such behaviour is consistent with our findings that bees can detect, and particularly learn to avoid, bitter surfaces like those in some pollen.

As far as we are aware, these data are the first to show that an insect can associate a bitter compound, here caffeine, or a salt detected at the gustatory periphery with a sucrose reward. Alkaloids like caffeine, and their taste can be associated with dopaminergic pathways and thus rewards in humans (Nehlig, 1999; Yeomans et al., 2007) and this may also have similar effects on bees (Wright et al., 2013). Aversions to peripheral stimulation by high levels of salt or bitter toxins, like the learnt responses of bees to quinine and caffeine surface chemistry, are more commonly reported in other insects (see above citations). For example, locusts use their tarsal sensilla to detect bitter compounds on plant surfaces prior to biting (Chapman and Ascoli-Christensen, 1999; White and Chapman, 1990). When they land on surfaces covered in alkaloids or salts, they move away (Rogers and Newland, 2000). Butterflies use tarsal chemoreception on leaves to identify host plants for oviposition; they also detect the compounds of non-host plants and avoid them (Ômura et al., 2011).

Previously, research in honeybees failed to find peripheral (antennal and tarsal) GRNs that spiked in response to stimulation with quinine (de Brito Sanchez et al., 2005; de Brito Sanchez et al., 2014). In the periphery and in the mouthparts, GRNs that respond to sugars reduce the rate of spiking when stimulated with bitter compounds like caffeine (de Brito Sanchez et al., 2005; de Brito Sanchez et al., 2014; Parkinson et al., 2023). In spite of our previous work that showed that honeybees have GRNs in the galea of the mouthparts that respond to quinine and amygdalin (Wright et al., 2010), some have argued that bees do not detect bitter *per se* but instead that bitter compounds ‘devalue’ the taste of nutrients like sucrose (Bestea et al., 2021; Lai et al., 2020). Our electrophysiological results, however, indicate that bumblebee species have the capacity to detect bitter compounds on their antennae using GRNs that spike in response to stimulation with quinine and caffeine. This the first report we know of where bitter sensitive GRNs have been identified in bumblebees and the first report that bee antennal GRNs sense bitter compounds. Our findings suggest either: bumblebees’ peripheral bitter taste capacity differs from that of honeybees, or that honeybees may similarly possess similar mechanisms that have not yet been discovered. The fact that there is a separate channel of information encoding the taste of bitter compounds or other ‘neutral’ tastants means that bees and other insects are likely to have the capacity to use this information to make independent associations of peripheral taste cues with unconditioned stimuli such as food rewards.

Bumblebees can learn and respond to several floral signalling modalities (see above), including those only detectable on contact with the flower surfaces. Such as, surface texture (Kevan and Lane, 1985; Whitney et al., 2009; Whitney et al., 2011) and temperature (Dyer et al., 2006; Harrap et al., 2017; Whitney et al., 2008). These can improve foraging decisions, even alongside other more conspicuous long-range cues (Goyret, 2010; Goyret and Raguso, 2006; Goyret et al., 2008; Harrap et al., 2020), enhancing foraging success (Raine and Chittka, 2007; Raine and Chittka, 2008). That bumblebees can utilise surface chemistry (peripheral taste) of artificial flowers to inform foraging choices suggests bees have, at least, the capacity to use surface chemistry as such a modality when visiting natural ‘real’ flowers. Whether petal surface chemistry is salient to bees or varies sufficiently for this to be advantageous is unclear. Flowers seem to have complex surface chemistry complements and petal surfaces differ in chemistry between flower species and from leaf surface chemistry (Giorio et al., 2015; Giorio et al., 2019; Goodwin et al., 2003; Griffiths et al., 2000; King et al., 2007; Shi et al., 2011; Tunstad et al., 2024). While petal surfaces are dominated by waxes, the presence of sugars and ‘bitter’ tastants (tannins and flavonoids) have been reported on the flower surface (Giorio et al., 2015; Giorio et al., 2019). The systemic nature of bitter toxins in plants that produce them (see above) means they are likely to occur on floral surfaces and vary in composition and occurrence in similar ways. This suggest bees might be able to distinguish different flowers, or floral and vegetive tissue, using surface chemistry. Bumblebees have been observed to avoid flowers producing high concentrations of alkaloids in nectar and plant tissues (Barlow et al., 2017; Wright et al., 2013), our results suggest associations between toxic rewards and bitter surface chemistry cues on the floral tissues may contribute to these responses. However, studies sampling the chemistry of the petal surface (epidermis, cuticle and epi-cuticle) are rare covering a handful of species (discussed in Tunstad et al., 2024). Furthermore, the artificial surface chemistry cues bees learnt in our experiments (comprising of plant-wood fibre and added tastants) differ from those of natural flowers. If present on flower surfaces, the specific tastants will be part of a complex mixture. Additionally, they may be imbedded within wax layers and not be accessible to the bee. Given bees possess both tarsal claws capable of imbedding in petals (Asperges et al., 2017; Pattrick et al., 2018) and dextrous antennae (Claverie et al., 2023; Ren et al., 2023), and interact with floral surfaces in complex ways (Barker et al., 2018; Biesmeijer et al., 2005; Hansen et al., 2012; Whitney et al., 2009), it is unclear what surface chemistry cues bees may have access to when visiting flowers. Regardless, our results do suggest flowers would benefit, in a manner similar to other modalities (Leonard et al., 2011b; Leonard et al., 2012), from having surface chemistry cues detectable to bees.

No spontaneous preferences toward surface chemistry were observed in naïve bees. Thus, our results may suggest bee responses to floral surface (petal, pollen or otherwise) chemistry are based on learning, not spontaneous preferences of naïve bees. This might be surprising, given the common instinctual feeding responses toward the same tastants in food (see citations above). However, responses to comparable stimuli between restrained and free-flying bees can vary (de Brito Sanchez et al., 2015; Mommaerts et al., 2013). Additionally, the tastants on the surface of our artificial flowers were dry and distinct from rewards bees consumed. This change in context may change how bees respond to the same stimuli (Leonard and Masek, 2014; Leonard et al., 2011b). It is also possible the surface chemistry stimuli were not intense enough to induce responses. Furthermore, the surface chemistry of artificial flowers used in testing likely differs from natural flowers (discussed above). Additionally, in nature the presence of these individual tastants may variably align with floral rewards; nectar varies in sugar content beyond just sucrose (Abrahamczyk et al., 2017; Parachnowitsch et al., 2019; Pattrick et al., 2024) and plants whose tissues contain salt and bitter compounds can have palatable and valuable nectar and pollen rewards to bees (Ayestaran et al., 2010; Barlow et al., 2017; Singaravelan et al., 2005; Tiedeken et al., 2014; Wright et al., 2013). These factors may mean, even though the tastants or similar chemicals, are encountered by bees on flower surfaces, bees have not been under selection to evolve instinctual preferences towards the surface chemistries presented in our experiments. In fact, in situations where a cue (across different flower species or time) does not always align with rewards, evolution of robust learning capacity toward that cue, like that bees show towards surface chemistry, is more adaptive (discussed in Dunlap et al., 2019).

Our results show peripheral taste capacity of bumblebees to be more complex than previously thought, with bumblebees being capable of using dry surface chemistry to inform flower foraging behaviour. Furthermore, we sound several ways bumblebee peripheral taste differs in mechanisms and responses compared to oral taste and taste of honeybees. Bumblebee peripheral taste able to directly detect bitter toxins and responses, at least appearing to be, dependent on learning. These findings expands our understanding of how bumblebees evaluate their food and the mechanisms of detecting toxicity by bees. Furthermore, identification of bumblebees’ abilities to use surface chemistry as a foraging cue warrants further exploration of the diversity of and responses of bees to natural floral and pollen surfaces, and nectar content, to assess how widely and in what contexts and to what extent this peripheral taste capacity and robust learning mechanisms associated with it are used in nature.

## Acknowledgements

We are grateful to Jen Scott, Rachel de Sousa and Jon Pattrick for technical support, assistance in the laboratory and their discussion of this work as it was being conducted.

## Author contributions

Conceptualization: G.A.W.; Methodology: M.J.M.H., R.H.P.; Formal analysis: M.J.M.H., R.H.P.; Investigation: M.J.M.H., R.H.P., H.J.; Resources: G.A.W.; Data curation: M.J.M.H., R.H.P.; Writing - original draft: M.J.M.H., R.H.P., G.A.W.; Writing - review & editing: M.J.M.H., R.H.P., H.J., G.A.W.; Visualization: M.J.M.H., R.H.P. Supervision: G.A.W.; Project administration: G.A.W.; Funding acquisition: G.A.W.

## Data availability

Raw data, data plotted in figures as well as code (where appropriate) necessary to generate graphical figures and repeat analysis are available at the following repositories. For artificial flower disc spectroscopy and SEM, and bee behavioural experiments (Harrap et al., 2025) - 10.6084/m9.figshare.27024322. For bee antennae SEM and electrophysiological recordings (Parkinson et al., 2025) - 10.6084/m9.figshare.28512926.

