## Supplementary material for "Bumblebees learn to use peripheral taste to predict the presence of nectar in flowers"

Pertaining to:

**A. Assessment of UV reflection of different flower variants**

All reflectance measurements were taken using an Ocean Optics Maya Pro 2000 spectrometer, a QR400-7-SR-BX probe and a DH-2000-BAL Deuterium Tungsten light source (all produced by Ocean Insight, Orlando, FL, USA). This spectrometer setup allowed measurement of reflection, expressed as a percentage of a standard. In all measurements reported a Spectralon standard (Labsphere, North Sutton, NH, USA) was used. The probe used measures reflectance across the UV-visible-IR spectra at wavelength intervals of approximately 0.5 nm (exact wavelengths of measurements vary).

Eight flowers of each flower variant (untreated, Sucrose, NaCl, Quinine and Caffeine flowers) were prepared (as described in the main text). A unique identifier number for each individual flower of each variant (untreated1, Sucrose1, etc.) was assigned to each and a this was marked by a sticker stuck to the side of the flower. Flowers were then taken to be measured for reflection in a sealed box (Really useful box 4 L, Shelving Plus, London) preventing acquisition of dust or contaminants that may alter reflection. Four reflectance spectra were then collected taking measurements about the flower top. Measurements were taken halfway between the feeding well and the edge of the flower top at 0, 90, 180 and 270 degrees (12 ,3, 6 and 9 o’clock positions) when viewing the flower such that the affixed marker sticker was facing the spectrometer operator.

While the spectrometer measures wavelengths across the bee’s sensitivity, reflectance measurements of paper between 420-700 nm were unreliable (relative to IR and UV ranges). This was due in part to the low illumination from the light source in this range and, thus, low reflectance from the standard in this range, and potentially florescence of the paper while under such high intensity UV light potentially causing artificial flowers to produce light in this range. This resulted in reflectance measurements in excess of 100% of the standard. Simulation of bee colour perception (Dyer and Chittka, 2004) was deemed not appropriate, due to unreliable estimates of reflection outside of the bee UV range. Flowers were not under intense UV light during trials, making any contribution of florescence to measured colour irrelevant for the purpose of assessment of visual differences between flower variants. Furthermore, all flower variants appeared white to humans, indicating an even reflection across these wavelengths. Assessment of potential for colour differences was therefore limited to the reflectance measurements in the 300-420 nm range, the bee-perceptible UV range. The bee’s capacity to distinguish flowers was then interpreted by bee’s capacity to distinguish similar UV reflection differences (e.g. Dyer and Chittka, 2004).

Reflectance spectra of different flower variants were then compared across the bee-perceptible UV range using a repeated measures ANOVA using R (R Core Team, 2022) and the LmerTest package (Kuznetsova et al., 2017). This compared how reflectance changed across wavelengths and how this varied between flower variants. Flower identity (40 flowers total) and specific measurement (160 specific measurements, i.e. each of the 4 measurements taken on each flower) were included as random factors that could influence overall reflectance (i.e. the ANOVA model intercept). More complex random effects were not included to facilitate model resolution and limit complexity. To allow ANOVA models to fit intercepts to reflectance at the lowest measured wavelength (300nm), and not 0nm, the effect of wavelength was represented by change in wavelength relative to the lowest wavelength measured (300.255 nm, with 300 nm having a value of 0 and 420 nm a value of 120 and so on). Data and code for spectra analyses can be found at Harrap et al. (2025).

Reflectance spectra of different flower variants were largely similar across the bee-perceptible UV range. Reflectance was generally low across the UV range but increased towards greater wavelengths, (Fig. S1). This is consistent with the flowers appearing white to humans, which would reflect more light at a consistent level across wavelengths greater than 400nm. These results, particularly the relatively low reflectance of flowers at the UV sensitivity peak of bees suggest all flower variants appear as a ‘green’ colour to bees, reflecting primarily a mix of human-blue and human-yellow wavelength light. Sucrose treated flowers, while showing similar shaped spectra, showed a slightly increased reflectance relative to untreated flowers (Fig. S1A). Quinine treated flowers had lower reflectance than untreated flowers between 300 and 340 nm (Fig. S1C). However, reflectance increased to slightly higher levels then untreated flowers at higher wavelengths. Caffeine (Fig. S1D) and NaCl (Fig. S1C) treated flower spectra remained like untreated flower spectra across the bee-perceptible UV range. Statistical analyses found spectra did differ between flower variants, as did the relationship between reflection and wavelength (table S1). However, assessment of ANOVA model fixed effects (table S1) revealed these differences were small and that NaCl treated flower spectra did not statistically differ from untreated flowers. The statistical difference found between flower variants by these analyses is likely influenced by the number of reflection measurements taken by the spectrometer (see wavelength relevant, degrees of freedom in table S1).

Taken together these findings indicate that, while evidence was found that flower variants (other than NaCl treated flowers) do not have identical bee-perceptible UV reflectance spectra, such spectra are similar across flower variants. The perceived colour of flowers by the bee will be dependent on both flower reflectance spectra and illumination while the bee encounters flowers. The detected differences in UV reflection are relatively small, with the general distribution and amount of reflectance being similar. These differences are unlikely to result in a perceptible difference in colour between flower variants that the bee could use to learn flower identity, especially when illuminated under less intense artificial lights in lab conditions, where UV illumination is lower and less spectrally diverse (see Dyer and Chittka, 2004).

**Figure S1:** The reflectance spectra of the different flower variant tops across bee UV receptor sensitive wavelengths. Reflectance is given as a percentage of the Spectralon standard at each wavelength. The solid black lines in each graph show the mean reflectance of untreated flowers at each wavelength (note: the untreated flower spectra are repeated across each panel to facilitate comparison). Solid blue lines indicate mean reflectance of A) Sucrose, B) NaCl, C) Quinine and D) Caffeine treated flowers across bee UV receptor sensitive wavelengths. Thin dashed lines above and below the mean spectra indicate standard error. Reflectance measurements are plotted against the left axis. Dotted green lines indicate the bee UV receptor sensitivity (measurement obtained from Dyer and Chittka, 2004) expressed as excitation at each wavelength as a percentage of the maximum excitation.

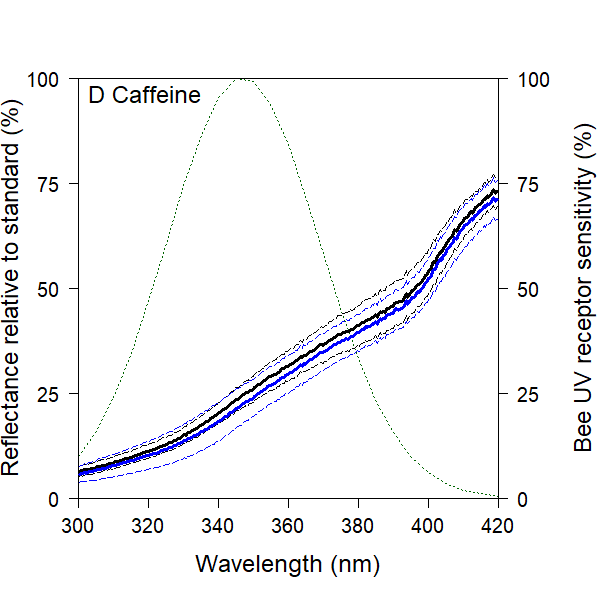

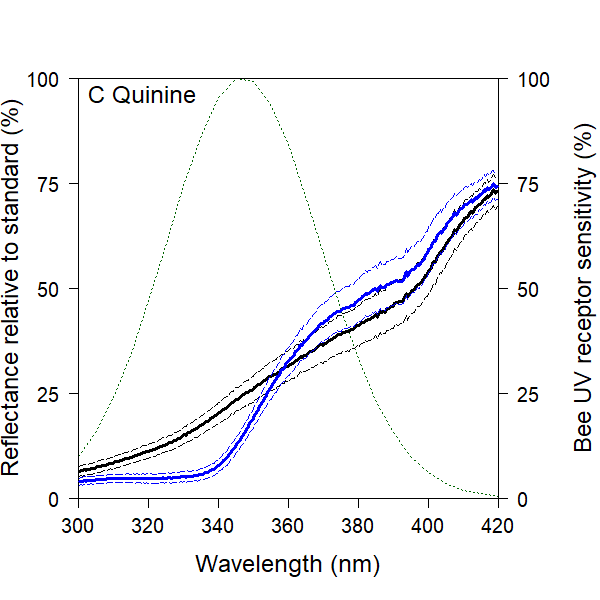

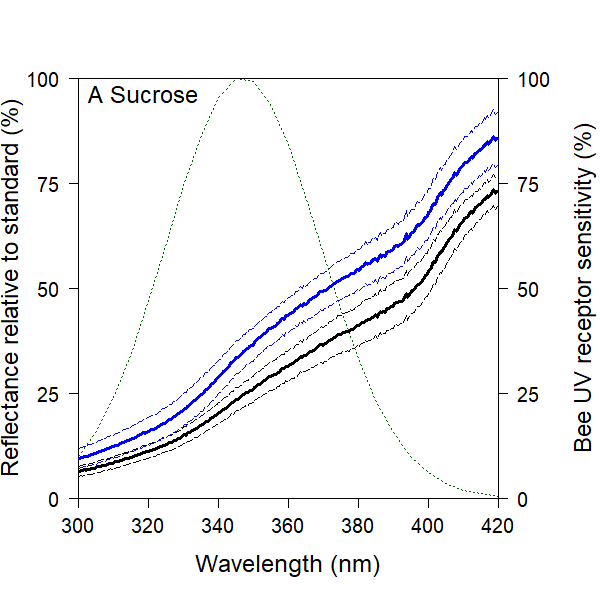

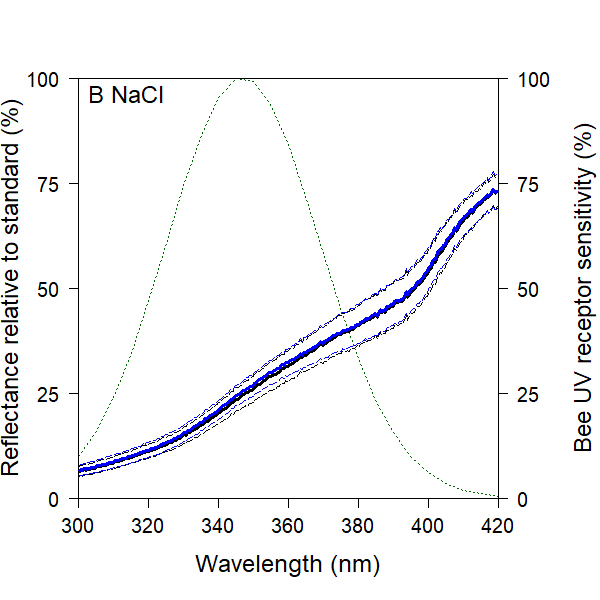

**Table S1:** Summary of the statistical analyses of flower variant spectra reflectivity (percent reflectance relative to a Spectralon standard at each wavelength) within the bee-perceptible UV range. Given are A) summary values for repeated measures ANOVA of flower spectra, and B) assessment of ANOVA model fixed effects. ‘*’ after P value indicates effect has a statistically significant influence on flower spectra (where *P*<0.05).

| **A) Repeated measures ANOVA** |  |  |  |  |  |
| --- | --- | --- | --- | --- | --- |
| **Effect** | **F** | **Nominator DF** | **Denominator DF** | **P** |  |
| Wavelength x Variant interaction | 2345.3 | 4 | 40960 | <0.001 | * |
| Variant | 59.3 | 4 | 41 | <0.001 | * |
| Wavelength | 970266.8 | 1 | 40960 | <0.001 | * |
| **B) Fixed effect assessment** |  |  |  |  |  |
| **Fixed Effect** | **estimate** | **t** | **DF** | **P** |  |
| Intercept | -0.07 | -0.12 | 41 | 0.91 |  |
| Wavelength (nm-300) | 0.55 | 407.52 | 40960 | <0.01 | * |
| Variant = Sucrose | 4.06 | 4.85 | 41 | <0.01 | * |
| Variant = NaCl | 0.50 | 0.56 | 41 | 0.58 |  |
| Variant = Quinine | -9.00 | -10.17 | 41 | <0.01 | * |
| Variant = Caffeine | -1.09 | -1.23 | 41 | 0.23 |  |
| Wavelength x Sucrose interaction | 0.10 | 52.48 | 40960 | <0.01 | * |
| Wavelength x NaCl interaction | <0.01 | 0.39 | 40960 | 0.70 |  |
| Wavelength x Quinine interaction | 0.13 | 68.11 | 40960 | <0.01 | * |
| Wavelength x Caffeine interaction | -0.01 | -4.34 | 40960 | <0.01 | * |

**B. Assessment of surface texture of different flower variants**

Small sections of each disc variant (untreated, sucrose, NaCl, Quinine and Caffeine) were imaged using the same SEM protocol applied to bee antennae, described in the main text. Three discs of each variate were imaged at 3 points randomly selected throughout each disc.

Example SEM images of each disk type at 500 times magnification are provided in Fig. S2. The full image dataset is available at Harrap et al. (2025). The surface texture of all disks variants was dominated by the random fibre structure of the filter paper disk themselves. Sucrose, quinine or caffeine treatments did not result in conspicuous changes in the disc surface structure relative to water treated discs. With no consistent difference in surface structure being visible at up to 500 times magnification (Fig. S2D). In discs treated with NaCl, small residue structures were visible across the paper fibres (Fig. S2D). However, these NaCl residues are irregularly arranged and smaller (below 5 µm) than surface structures bees are typically able detect (Kevan and Lane, 1985; Whitney et al., 2009; Whitney et al., 2011). Based on SEM imaging, chemical treatments did not alter the surface texture in a way that bees could detect.

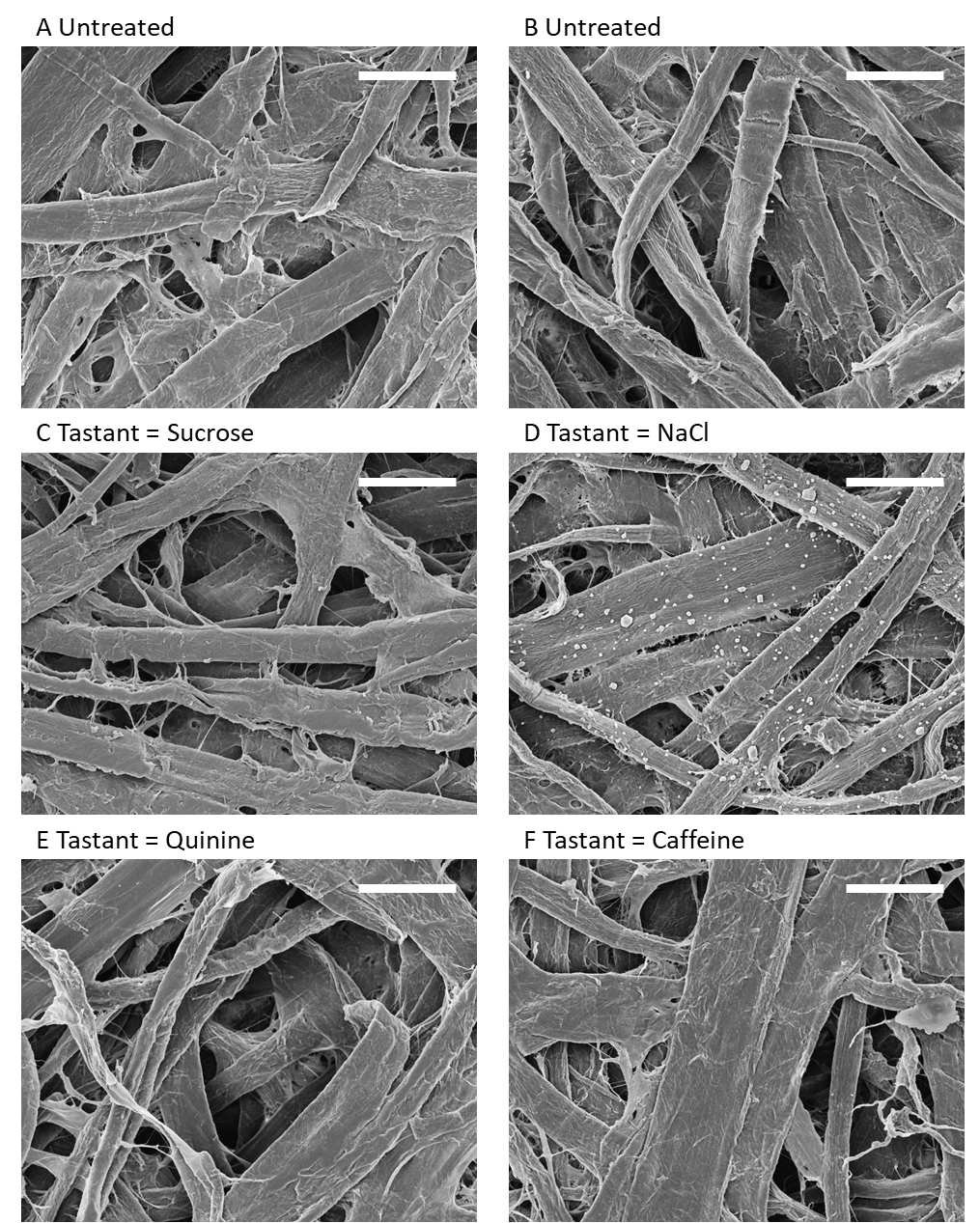
**Figure S2:** SEM images of artificial flower discs at x500 magnification. Treatment applied to discs imaged is as follows: A and B) water treated control discs, C) Sucrose discs, D) NaCl discs, E) Quinine discs and F) Caffeine discs. White scalebar in top right of each image represents 50 µm in all panels. The full image dataset is available at Harrap et al. (2025).

**C. Statistical models and simplification procedure for bee training phase of differential conditioning experiments**

The following represents the initial model of bee learning (before any simplification was applied) fit to the training phase responses of bees in the control and the tastant positive and negative conditioning groups of each tastant:

$y_{nx}=i+\left( v \times l \right)+P\left( s_{p}+\left( v \times c_{p} \right) \right)+M\left( s_{m}+\left( v \times c_{m} \right) \right)+ b_{n}+(v \times r_{n})$.	(1)

Where $y_{nx}$is the arcsine square-root success rate of bee $n$ over the previous 10 visits to the artificial flowers, at $x$ flower visits. $v$ relates to the number of flower visits the bee has made to the artificial flowers, $x$, by the following:

$v=x-10$.	(2)

The data for $y$ are calculated in blocks of 10 visits (i.e. at 10, 20, 30, 40, 50, 60 and 70 visits). The transformation shown in Eqn 2 allows the model intercept to be the first calculated success rate (that achieved by bees at 10 visits). Note that, due to the linear shape of success-experience relationships, unlike previous studies that used similar models (e.g. Harrap et al., 2017; Harrap et al., 2021), $x$ was not log-transformed. Parameter $i$ is the initial arcsine square-root success rate, the model intercept, for bees in the control group when $x=10$. Parameter $l$ dictates the change in arcsine square-root success rate with increased $x$ in the control group, thus $l$ is effectively the ‘learning speed’ parameter and allows the bee's experience to effect success rate. $P$ and $M$ are Boolean parameters that allow the model to alter $y$ if the bee is not in the control group. Parameter $P$ indicates whether bee $n$ is in the tastant positive group, where:

$P= \left\{ \begin{aligned} 0 \\ 1 \end{aligned} \right. \begin{matrix} bee is not in the tastant positive group \\ bee is in the tastant positive group \end{matrix}$ ,	(3)

while Boolean $M$ whether bee $n$ is in the tastant negative group

$M= \left\{ \begin{aligned} 0 \\ 1 \end{aligned} \right. \begin{matrix} bee is not in the tastant negative group \\ bee is in the tastant negative group \end{matrix}$ .	(4)

Parameters $s_{p}$ and $c_{p}$ are the change in initial success rate and learning speed, relative to $i$ and $l$ respectively of bees in the tastant positive conditioning group. Likewise, Parameters $s_{m}$ and $c_{m}$ are the change in initial success rate and learning speed, relative to $i$ and $l$, of bees in the tastant negative conditioning group. Variation between individual bees was included in our model as a random factor. Parameters $b_{n}$ and $r_{n}$ represent the change in initial arcsine success rate and learning speed, for bee number $n$, relative to $i$ and $l$. In the model described in Eqn 1 parameters $i$, $l$, $s_{p}$, $c_{p}$, $s_{m}$, $c_{m}$, $b_{n}$and $r_{n}$are parameters to be estimated.

The model simplification procedure involved paired comparisons between the standing ‘best model’, beginning with the full model described in Eqn 1, with a simpler model. Simpler models were constructed from the standing best model but with further parameters removed (effectively forcing the relevant parameters to equal zero) and tested against the standing best model in the sequence described below. Should the simpler model have a lower AIC or be comparable to the standing best fitting model based on AIC, as laid out by Richards (2008), this simpler model would become the standing best model for the next comparison. If removal of a parameter led to a significant increase in AIC, the standing best (more complex) model would remain the best for the next comparison.

For each tastant treatment, the influence of random factors are compared first. This is done via comparison of the initial model with one without $r_{n}$. This comparison tests if individual bees differed only in intercepts or intercepts and learning speed (as in the initial model). Next, we tested for the presence of interacting effects between conditioning group and experience, that is individual learning speeds for each conditioning group. This was done by comparing the best model with one where $c_{p}$ and $c_{m}$ were removed. Removal of these parameters would create a model where the relationship between $y$ and $x$ was determined only by a common learning speed, $l$, across conditioning groups. The next comparison compared models without this common learning speed with the standing best model. This tested whether groups had a common learning speed, if previous comparisons indicated the standing best model should not include $c_{p}$ and $c_{m}$. Alternatively, this comparison tested for the presence of a baseline improvement in success in control group bees the standing best model included $c_{p}$ and $c_{m}$. Lastly, models where conditioning groups had a shared intercept (dictated by $i$) were compared to the standing best model by removing parameters $s_{p}$ and $s_{m}$. This tested whether bees in different conditioning groups differed in success rate, independent of changes in success with experience. The standing best model at the end of this comparison was considered the most parsimonious model of training phase responses for that tastant overall.

To assess between conditioning group differences more directly, a further set of models were fit to data of paired conditioning groups. These models took the same structure as the model given in Eqn 1 but were fit to the training phase response data of: the control group and the tastant positive group; the control group and the tastant negative group; and the tastant positive and negative group, of each tastant (3 sets of models per tastant). If the previous comparisons for that tastant indicated that parameters $l$, $c_{p}$ and $c_{m}$ or $b_{n}$ should not be in the most parsimonious model they were excluded, as they were already established to have no effect. These models were compared to identical models where parameters $c_{p}$ and $c_{m}$ were removed (if they were included at all), and then a model where $s_{p}$ and $s_{m}$ were removed. The remaining parameters in the final of these paired group models, indicated whether the paired groups differed in slope (if $c_{p}$ and $c_{m}$ were retained) or intercept (where only $s_{p}$ and $s_{m}$ were).

**Table S2:** The results of model selection for bee training phase responses in the sucrose conditioning groups (sucrose positive, ‘+ve’, and sucrose negative, ‘-ve’) and the control group (‘Con.’). Comparisons of standing best models fit to all three groups and a simpler models where each parameter is removed in turn are given. As are the results of between group comparisons where effects are tested on models fit to data of paired conditioning groups; the shaded, right aligned entries. At each comparison AIC is given for both models as is an assessment of model fit (where Δdeviance = ‘Δdev’). Bold AIC value indicates the best model from this comparison, according to Richards (2008). Note that here a ΔAIC of at least 6 is needed for a more complex model to be favoured in a comparison. A verdict on each comparison is given: Asterisks ‘✱’ indicate parameters where models including them perform better in terms of AIC and are thus included in the most parsimonious model. Daggers ‘†’ indicate parameters where models including them do not perform better in terms of AIC but fit is still significantly improved. Blank values of ‘verdict’ indicate parameters were neither AIC performance of fit are improved when including them in models.

| Tastant = Sucrose | | | | | | | | | |
| --- | --- | --- | --- | --- | --- | --- | --- | --- | --- |
| Step | Focal Parameter/ tested effect |  | Standing best model AIC | Simpler model  AIC | ΔAIC | Δdev. | df | *p* | Verdict |
| 1 | Random slopes | $r_{n}$ | -241.24 | **-244.17** | 2.93 | 1.08 | 2 | 0.584 |  |
| 2 | Interaction effects | $c$ | -244.17 | **-241.06** | 3.11 | 7.11 | 2 | 0.029 | † |
| 3 | Learning | $l$ | **-241.06** | -197.72 | 43.34 | 45.34 | 1 | <0.001 | 🞷 |
| 4 | Differing Intercepts | $s$ | **-241.06** | -224.45 | 16.61 | 20.61 | 2 | <0.001 | 🞷 |
| 4a | Distinction between  Con. & +ve | $s$ | **-162.05** | -153.15 | 8.9 | 10.90 | 1 | 0.001 | 🞷 |
| 4b | Distinction between  Con. & -ve | $s$ | **-163.21** | -142.60 | 20.61 | 22.61 | 1 | <0.001 | 🞷 |
| 4c | Distinction between  +ve & -ve | $s$ | -157.56 | **-159.03** | 1.47 | 0.54 | 1 | 0.464 |  |

**Table S3:** The results of model selection for bee training phase responses in the NaCl conditioning groups (NaCl positive, ‘+ve’, and NaCl negative, ‘-ve’) and the control group (‘Con.’). See table S2 for detail on reported values within the table.

| Tastant = NaCl | | | | | | | | | |
| --- | --- | --- | --- | --- | --- | --- | --- | --- | --- |
| Step | Tested effect |  | Standing best model AIC | Simpler model  AIC | ΔAIC | Δdev. | df | *p* | Verdict |
| 1 | Random slopes | $r_{n}$ | -229.44 | **-227.88** | 1.56 | 5.56 | 2 | 0.062 |  |
| 2 | Interaction effects | $c$ | -227.88 | **-230.72** | 2.84 | 1.16 | 2 | 0.561 |  |
| 3 | Learning | $l$ | **-230.72** | -203.37 | 27.35 | 29.35 | 1 | <0.001 | 🞷 |
| 4 | Differing Intercepts | $s$ | **-230.72** | -224.08 | 6.64 | 10.64 | 2 | 0.005 | 🞷 |
| 4a | Distinction between  Con. & +ve | $s$ | -145.87 | **-140.71** | 5.16 | 7.16 | 1 | 0.007 | † |
| 4b | Distinction between  Con. & -ve | $s$ | **-167.28** | -159.93 | 7.35 | 9.35 | 1 | 0.002 | 🞷 |
| 4c | Distinction between  +ve & -ve | $s$ | -144.42 | **-146.40** | 1.98 | 0.02 | 1 | 0.889 |  |

**Table S4:** The results of model selection for bee training phase responses in the quinine conditioning groups (quinine positive, ‘+ve’, and quinine negative, ‘-ve’) and the control group (‘Con.’). See table S2 for detail on reported values within the table.

| Tastant = Quinine | | | | | | | | | |
| --- | --- | --- | --- | --- | --- | --- | --- | --- | --- |
| Step | Tested effect |  | Standing best model AIC | Simpler model  AIC | ΔAIC | Δdev. | df | *p* | Verdict |
| 1 | Random slopes | $r_{n}$ | -223.79 | **-225.43** | 1.64 | 2.37 | 2 | 0.306 |  |
| 2 | Interaction effects | $c$ | -225.43 | **-227.30** | 1.87 | 2.13 | 2 | 0.346 |  |
| 3 | Learning | $l$ | **-227.30** | -211.59 | 15.71 | 17.70 | 1 | <0.001 | 🞷 |
| 4 | Differing Intercepts | $s$ | **-277.30** | -220.92 | 6.38 | 10.36 | 2 | 0.006 | 🞷 |
| 4a | Distinction between  Con. & +ve | $s$ | **-176.82** | -169.99 | 6.83 | 8.83 | 1 | 0.003 | 🞷 |
| 4b | Distinction between  Con. & -ve | $s$ | -136.43 | **-130.67** | 5.76 | 7.75 | 1 | 0.005 | † |
| 4c | Distinction between  +ve & -ve | $s$ | -140.93 | **-142.44** | 1.51 | 0.48 | 1 | 0.487 |  |

**Table S5:** The results of model selection for bee training phase responses in the caffeine conditioning groups (caffeine positive, ‘+ve’, and caffeine negative, ‘-ve’) and the control group. See table S2 for detail on reported values within the table.

| Tastant = Caffeine | | | | | | | | | |
| --- | --- | --- | --- | --- | --- | --- | --- | --- | --- |
| Step | Tested effect |  | Standing best model AIC | Simpler model  AIC | ΔAIC | Δdev. | df | *p* | Verdict |
| 1 | Random slopes | $r_{n}$ | -229.07 | **-231.65** | 2.58 | 1.42 | 2 | 0.491 |  |
| 2 | Interaction effects | $c$ | -231.65 | **-234.78** | 3.13 | 0.86 | 2 | 0.650 |  |
| 3 | Learning | $l$ | **-234.78** | -212.70 | 22.08 | 24.09 | 1 | <0.001 | 🞷 |
| 4 | Differing Intercepts | $s$ | -234.78 | **-231.34** | 3.44 | 7.45 | 2 | 0.024 | † |
| 4a | Distinction between  Con. & +ve | $s$ | -164.90 | **-160.47** | 4.43 | 6.43 | 1 | 0.011 | † |
| 4b | Distinction between  Con. & -ve | $s$ | -152.18 | **-149.26** | 2.92 | 4.92 | 1 | 0.027 | † |
| 4c | Distinction between  +ve & -ve | $s$ | -147.58 | **-149.57** | 1.99 | 0.013 | 1 | 0.909 |  |
